# Differential methylation between ethnic sub-groups reflects the effect of genetic ancestry and environmental exposures

**DOI:** 10.1101/036822

**Authors:** Joshua M. Galanter, Christopher R. Gignoux, Sam S. Oh, Dara Torgerson, Maria Pino-Yanes, Neeta Thakur, Celeste Eng, Donglei Hu, Scott Huntsmann, Harold J. Farber, Pedro C Avila, Emerita Brigino-Buenaventura, Michael A LeNoir, Kelly Meade, Denise Serebrisky, William Rodríguez-Cintrón, Rajesh Kumar, Jose R Rodríguez-Santana, Max A. Seibold, Luisa N. Borrell, Esteban G. Burchard, Noah Zaitlen

**Author notes:** These authors contributed equally to this work. Please address correspondence to: Esteban G. Burchard, MD, MPH, University of California, San Francisco, Departments of Bioengineering & Therapeutic Sciences and Medicine UCSF Box 2911, San Francisco, CA 94143-2911, Noah Zaitlen, PhD, University of California, San Francisco Department of Medicine, UCSF Box 2552, San Francisco, CA 94143-2552, Joshua Galanter, MD, MAS, University of California, San Francisco, Departments of Medicine and Epidemiology & Biostatistics UCSF Box 2911, San Francisco, CA 94143-2552.

## Abstract

In clinical practice and biomedical research populations are often divided categorically into distinct racial/ethnic groups. In reality, these categories, which are based on social rather than biological constructs, comprise diverse groups with highly heterogeneous histories, cultures, traditions, religions, social and environmental exposures and ancestral backgrounds. Their use is thus widely debated and genetic ancestry has been suggested as a complement or alternative to this categorization. However, few studies have examined the relative contributions of racial/ethnic identity, genetic ancestry, and environmental exposures on well-established and fundamental biological processes. We examined the associations between ethnicity, ancestry, and environmental exposures and DNA methylation. We typed over 450,000 CpG sites in primary whole blood of 573 individuals of diverse Hispanic descent who also had high-density genotype data. We found that both self-identified ethnicity and genetically determined ancestry were significantly associated with methylation levels at a large number of CpG sites (916 and 194, respectively). Among loci differentially methylated between ethnic groups, a median of 75.7% (IQR 45.8% to 92%) of the variance in methylation associated with ethnicity could be accounted for by shared genomic ancestry accounts. We also found significant enrichment (p = 4.2 × 10^-64^) of ethnicity-associated sites amongst loci previously associated with environmental and social exposures, particularly maternal smoking during pregnancy. Our study suggests that although differential methylation between ethnic groups can be partially explained by the shared genetic ancestry, a significant effect of ethnicity is likely due to environmental, social, or cultural factors, which differ between ethnic groups.

**One Sentence Summary:** In order to better understand the role of ethnic self-identification and genetically determined ancestry in biomedical outcomes, we explore their relative contributions to variation in methylation, a fundamental biological process.

**Sources of Funding:** This research was supported in part by the Sandler Family Foundation, the American Asthma Foundation, National Institutes of Health (P60 MD006902, R01 HL117004, R21ES24844, U54MD009523, R01 ES015794, R01 HL088133, M01 RR000083, R01 HL078885, R01 HL104608, U19 AI077439, M01 RR00188, U01 HG009080, and R01 HL135156), ARRA grant RC2 HL101651, and TRDRP 24RT-0025; EGB was supported in part through grants from the Flight Attendant Medical Research Institute (FAMRI), and NIH (K23 HL004464); NZ was supported in part by an NIH career development award from the NHLBI (K25HL121295). JMG was supported in part by NIH Training Grant T32 (T32GM007546) and career development awards from the NHLBI (K23HL111636) and NCATS (KL2TR000143) as well as the Hewett Fellowship; N.T. was supported in part by an institutional training grant from the NIGMS (T32-GM007546) and career development awards from the NHLBI (K12-HL119997 and K23-HL125551), Parker B. Francis Fellowship Program, and the American Thoracic Society; CRG was supported in part by NIH Training Grant T32 (GM007175) and the UCSF Chancellor’s Research Fellowship and Dissertation Year Fellowship; RK was supported with a career development award from the NHLBI (K23HL093023); HJF was supported in part by the GCRC (RR00188); PCA was supported in part by the Ernest S. Bazley Grant; MAS was supported in part by 1R01HL128439-01. This publication was supported by various institutes within the National Institutes of Health. Its contents are solely the responsibility of the authors and do not necessarily represent the official views of the NIH.

## Introduction

Race, ethnicity, and genetic ancestry have had a complex and often controversial history within biomedical research and clinical practice [1–3]. For example, race- and ethnicity-specific clinical reference standards are based on population-based sampling on a given physical trait such as pulmonary function[4,5]. However, because race and ethnicity are social constructs and poor markers for genetic diversity, they fail to capture the heterogeneity present within racial/ethnic groups and in admixed populations[6]. To account for these heterogeneities and to avoid social and political controversies, the genetics community has grouped individuals by genetic ancestry instead of race and ethnicity[3]. Indeed, recent work from our group and others have demonstrated that genetic ancestry improves diagnostic precision compared to racial/ethnic categorizations for specific medical conditions and clinical decisions[7–9].

However, racial and ethnic categories also reflect the shared experiences and exposures to known risk factors for disease, such as air pollution and tobacco smoke, poverty, and inadequate access to medical services, which have all contributed to worse disease outcomes in certain populations[10,11]. Thus, it is unclear whether defining groups through genetic ancestry can capture these shared exposures. In this work we seek to explore the relative contributions of genetically defined ancestry and social, cultural and environmental factors to understanding differential methylation between ethnic groups.

Epigenetic modification of the genome through methylation plays a key role in the regulation of diverse cellular processes[12]. Changes in DNA methylation patterns have been associated with complex diseases, including various cancers[13], cardiovascular disease[14,15], obesity[16], diabetes[17], autoimmune and inflammatory diseases[18], and neurodegenerative diseases[19]. Epigenetic changes are thought to reflect influences of both genetic[20] and environmental factors[21], and have been shown to vary between racial groups[22]. The discovery of methylation quantitative trait loci (meQTL’s) across populations by Bell et al. established the influence of genetic factors on methylation levels in a variety of tissue types[20], with meQTL’s explaining between 22% and 63% of the variance in methylation levels. Multiple environmental factors have also been shown to affect methylation levels, including endocrine disruptors, tobacco smoke[23,24], polycyclic aromatic hydrocarbons, infectious pathogens, particulate matter, diesel exhaust particles[25], allergens, heavy metals, and other indoor and outdoor pollutants[26]. Psychosocial factors, including measures of traumatic experiences[27–29], socioeconomic status[30,31], and general perceived stress[32], also affect methylation levels. Since both genetic and environmental exposures affect methylation, this represents an ideal phenotype to explore the relative contributions of these two factors on differential methylation between ethnic groups.

In this work, we leveraged genome-wide methylation data in 573 Latino children of diverse Latino sub-ethnicities enrolled in the Genes-Environment and Admixture in Latino Americans (GALA II) study[33] whose genetic ancestry had been determined from dense genotyping arrays. This allowed us to explore the extent to which the differences in methylation between Latino sub-groups could be explained by their shared genetic ancestry. We found that many of the methylation differences associated with ethnicity could be explained by shared genetic ancestry. However, even after adjusting for ancestry, significant differences in methylation remained between the groups remained at multiple loci, reflecting social and environmental influences upon methylation.

Our findings have important implications for both the use of ancestry to capture biological changes and of race/ethnicity to account for social and environmental exposures. Epigenome-wide association studies in diverse populations may be susceptible to confounding due to environmental exposures in addition to confounding due to population stratification[34]. The findings also have implications for the common practice of considering individuals of Latino descent, regardless of origin as a single ethnic group.

## Results

The study included 573 participants, the majority of whom self-identified as being either of Puerto Rican (n = 220) or Mexican origin (n = 276). Table 1 displays baseline characteristics of the GALA II study participants with methylation data included in this study, stratified by ethnic subgroups (Puerto Rican, Mexican, Other Latino, and Mixed Latinos who had grandparents of more than one national origin). Figure S1 shows the distribution of African, European, and Native American ancestry among the 524 participants with genomic ancestry estimates.

**TABLE 1:**
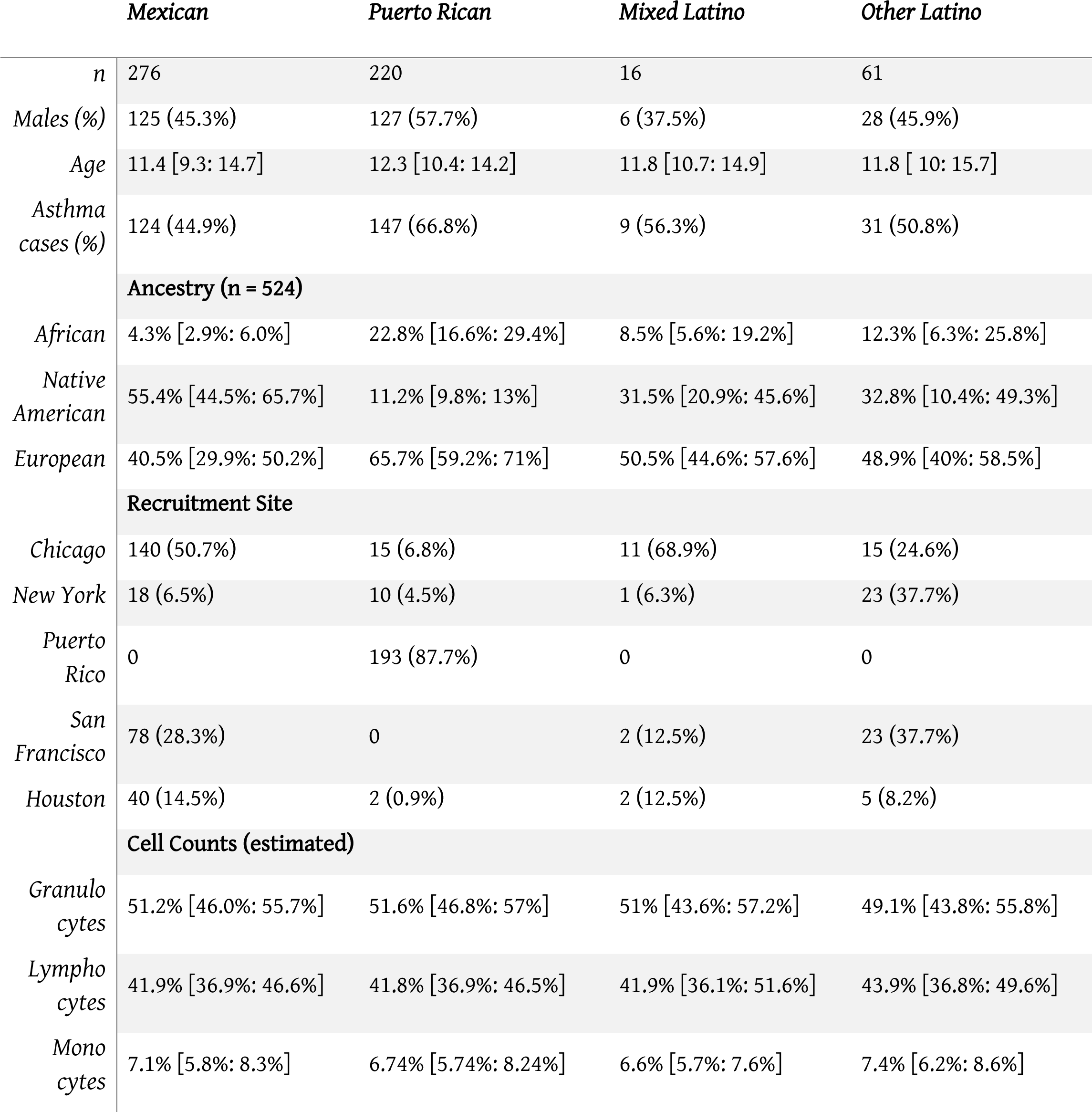
Baseline characteristics of GALA II participants with methylation data, stratified by ethnicity.

### Global patterns of methylation

Differences in ethnicity and ancestry resulted in discernible patterns in the global methylation profile as demonstrated in a multidimensional scaling analysis [Figure S2A]. As expected[30,35], the first few principal coordinates are strongly correlated to imputed cell composition [Figure S2B-C]. There are also significant associations of self-identified sub-ethnicity with PC2 (p-ANOVA = 0.003), PC3 (p-ANOVA = 0.004), PC6 (p-ANOVA = 0.0001), PC7 (p-ANOVA = 0.0003) [Figure S3A], and PC8 (p-ANOVA = 0.0003), after adjusting for age, sex, disease status, cell components, and technical factors (plate and position). Genetic ancestry was associated with PC3 (p-ANOVA = 0.002), PC7 (p-ANOVA = 0.0004) [Figure S3B] and PC8 (p-ANOVA = 0.001) in a two degree of freedom ANOVA test, adjusting for age, sex, disease status, cell components, technical factors, and ethnicity. Table S1 summarizes the results of the simple correlation analysis of methylation with ethnicity and ancestry, as well as the adjusted nested ANOVA models described above and the mediation results described below.

A mediation analysis[36] revealed that the associations between ethnicity and PCs 3, 7, and 8 were significantly mediated by Native American ancestry (mediation p = 0.01, <0.001, and <0.001, respectively) and inclusion of Native American ancestry in the regression model of PCs 3, 7, and 8 caused the ethnicity associations to be non-significant. However, the associations of ethnicity with PCs 2 and 6 were not explained by Native American, African or European ancestry (mediation p > 0.05), suggesting that the ethnic differences in these principal components are associated with global methylation patterns not captured by the shared genetic ancestry of each ethnic group. When genetic ancestry was regressed on the methylation data with the principal coordinates recalculated using the residuals of the regression between methylation and ancestry, there was an association between ethnicity and PC6 (p-ANOVA = 0.003). However, there was no association with any of the other principal coordinates. These observations suggest that while shared genetic ancestry can explain some of the association between ethnicity and global methylation patterns, other non-genetic factors, such as environmental and social exposure differences associated with ethnicity influence methylation and are not captured by measures of genetic ancestry.

### Epigenome-wide association of self-identified ethnicity

An epigenome-wide association study of self-identified ethnicity (see methods for details of ascertainment of ethnicity) and methylation identified a significant difference in methylation M-values between ethnic groups at 916 CpG sites at a Bonferroni-corrected significance level of less than 1.6×10^-7^ [Figure 1A and Table S2]. The most significant association with ethnicity occurred at cg12321355 in the ABO blood group gene (*ABO*) on chromosome 3 (p-ANOVA 6.7×10^-22^) [Figure 1B]. A two degree of freedom ANOVA test for genomic ancestry was also significantly associated with methylation level at this site (p = 2.3×10^-5^) [Figure 1C], and when the analysis was stratified by ethnic sub-group, showed an association in both Puerto Ricans and Mexicans (p = 0.001 for Puerto Ricans, p = 0.003 for Mexicans). Although adjusting for genomic ancestry attenuated the effect of ethnicity, a significant association between ethnicity and methylation remained (p = 0.04). Recruitment site, an environmental exposure proxy, was not significantly associated with methylation at this locus (p = 0.5), suggesting that environmental differences associated with ethnicity beyond geography and ancestry are driving the association.

**Figure 1:**
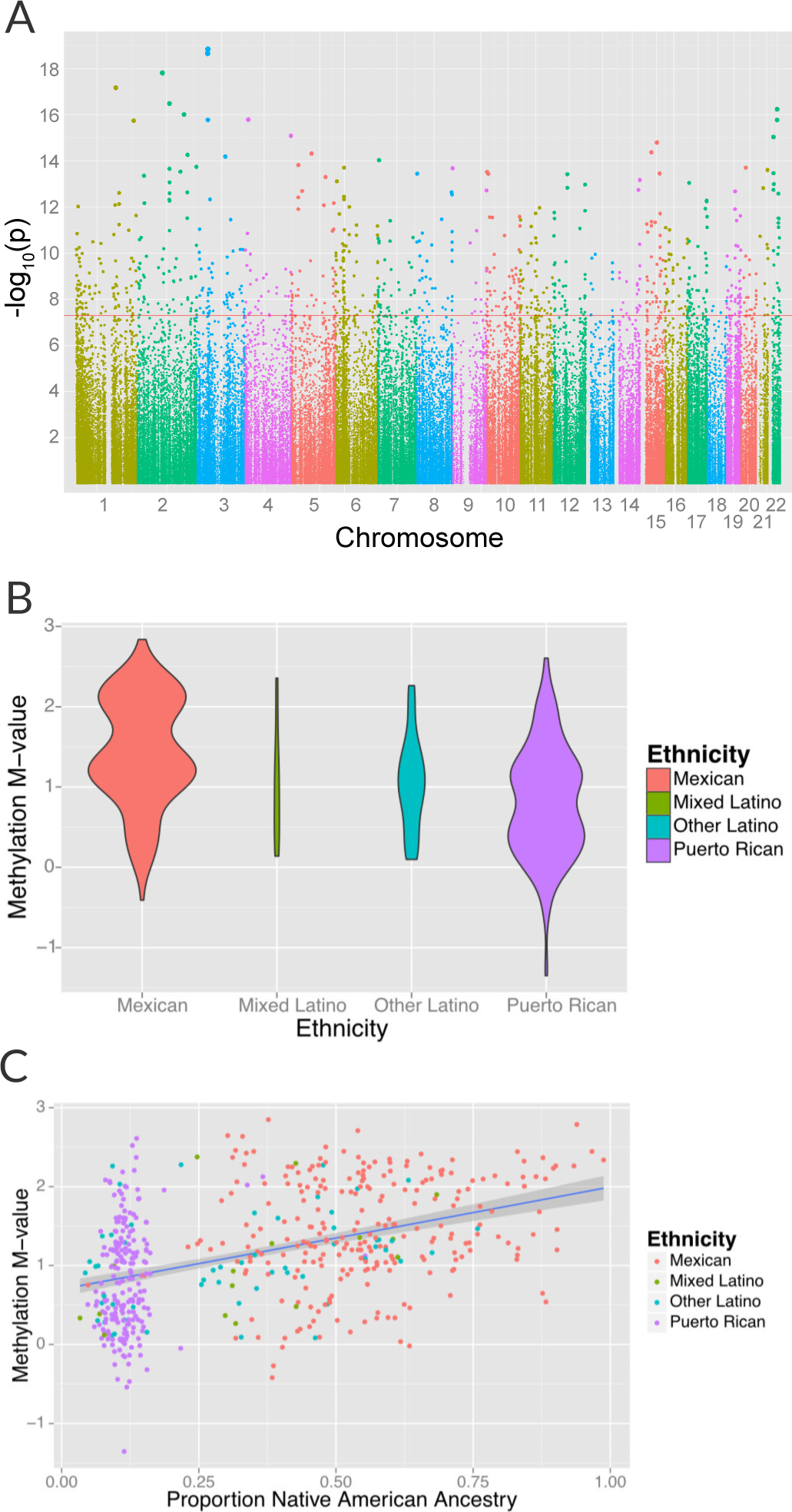
Associations between ethnicity and methylation (**A**) Manhattan plot showing the associations between ethnicity and methylation at individual CpG loci. (**B**) Violin plot showing one such locus, cg19145607. Mexicans are relatively hypermethylated compared to Puerto Ricans (p = 1.4×10^-19^). (**C**) Plot showing the association between Native American ancestry at the locus and methylation levels at the locus colored by ethnicity; Native American ancestry accounts for 58% of the association between ethnicity and methylation at the locus.

In order to determine the relative contribution of shared genetic ancestry and other factors associated with ethnicity, we repeated the analysis adjusting for ancestry. A significant association remained in 314 of the 834 (37.8%, p = 1.7 × 10^-183^ for enrichment) CpG sites associated with ethnicity [Figure S4A and Table S2] (82 sites were excluded because they demonstrated unstable coefficient estimates and inflated standard errors due to strong correlations between ethnicity and ancestry, especially Native American ancestry [see Figure S1]). Genomic ancestry explained a median of 4.2% (IQR 1.8% to 8.3%) of the variance in methylation at these loci and accounts for a median of 75.7% (IQR 45.8% to 92%) of the total variance in methylation explained jointly by ethnicity and ancestry [Figure S4B]. Sensitivity tests for departures from linearity, fine scale population substructure and the exclusion of the 16 participants who self-identified as “Mixed Latino” sub-ethnicity, did not meaningfully affect our results [See Supplementary Text and Tables S2 – S6]. To rule out any residual confounding due to recruitment sites, we conducted an additional analysis on the effect of recruitment site on methylation both for the overall study and for the Mexican participants (the largest study population in this analysis). We observed no significant independent effect of recruitment site suggesting that confounding due to recruitment region was limited, at least within the United States.

To explore the effect of departures from a linear association between ancestry and methylation, we incorporated both higher order polynomials and cubic splines of ancestry into our models. We observed a significant departure from linearity (p < 0.05) in only 26 (for splines) and 25 (for polynomials) of the 314 CpG’s where an association between ethnicity and methylation remained after adjusting for ancestry; however, the association between ethnicity and methylation remained even after adjusting for non-linearity at all sites [Tables S3 and S4].

Environmental differences between geographic locations or recruitment sites are a potential non-genetic explanation for ethnic differences in methylation. We investigated the independent effect of recruitment site on methylation by analyzing the associations between recruitment site and individual methylation loci after adjusting for ethnicity. We did not find any loci significantly associated with recruitment site at a significance threshold of 1.6 × 10^-7^. We then performed an analysis to assess the effect of recruitment sites on methylation stratified by ethnicity. We did not find any loci significantly associated with recruitment site and methylation among Mexican participants. We were underpowered to perform a similar analysis for Puerto Ricans because there were only 27 Puerto Rican participants recruited outside of Puerto Rico. To ensure that the absence of association in Mexicans was not due to the loss of power from the smaller sample size, we repeated our analysis of the association between ethnicity and ancestry randomly down-sampling to 276 participants to match the sample size in the analysis of geography in Mexicans. While down-sampling the study to this degree resulted in a loss of power, 128 methylation sites were still associated with ancestry. We conclude that recruitment site was unlikely to be a significant confounder of our associations between ethnicity and methylation and was not a significant independent predictor of methylation.

While most population substructure in Latinos would be expected to arise from differences in continental ancestry[37,38], there is evidence of finer scale (sub-continental) ancestry in Latino populations[39]. We tested for the effect of fine scale substructure by calculating principal components for all participants with genotyping data using Eigensoft[40]. We found significant associations between principal components 3-10 (PC’s 1 and 2 were almost perfectly collinear with ancestry, with an adjusted R^2^ > 0.998 for all three ancestry proportions, and were therefore excluded) and ethnicity. We therefore added these 8 PC’s to models of ethnicity and methylation, and found an association between these genetic PC’s and methylation in 63/314 CpG’s that had remained associated with ethnicity after adjusting for ancestry. Adjusting for higher order substructure in these CpG’s explained the association between ethnicity and methylation in 51 additional loci. This left 263 loci associated with ethnicity after adjustment for ancestry where there was either no association between PC’s 3-10 and methylation or the inclusion of these PC’s did not affect the association between ethnicity and methylation.[Table S5]

As only 16 participants self-identified as “Mixed Latino”, we performed a sensitivity analysis to test the effect of excluding these participants from the analysis and only examining Puerto Ricans, Mexicans, and “Other Latinos”. We found that excluding self-identified “Mixed Latino” participants from the analysis did not significantly alter the results in most cases [Table S6]. Of the 916 CpG’s associated with ethnicity at a genome-wide scale (p < 1.6 × 10-7) in models including individuals self-identified as “Mixed Ethnicity”, 894 (97.5%) were still significant at a genome-wide scale when “Mixed Latinos” were excluded. All but two of the CpG’s that did not meet genome-wide significance were significant when correcting for 916 tests (p < 5×10-5). In addition, an additional 290 CpG loci that did not meet genome-wide significance in the original analysis were significant at a genome-wide scale when self-identified “Mixed Latinos” were excluded. While these loci did not meet genome-wide significance in the original analysis that included Mixed Latinos, they all had p-values lower than 2 ×10^-6^. Thus we conclude that a sensitivity test excluding individuals of mixed Latino ethnicity did not significantly alter the conclusions.

We conclude that shared genetic ancestry explains much but not all of the association between ethnicity and methylation. Other, non-genetic factors associated with ethnicity likely explain the ethnicity-associated methylation changes that cannot be accounted for by genomic ancestry alone.

### Ethnic differences in environmentally-associated methylation sites

Methylation at CpG loci that had previously been reported to be associated with environmental exposures whose exposure prevalence differs between ethnic groups were tested for association with ethnicity in this study. A recent meta-analysis of maternal smoking during pregnancy, an exposure that varies significantly by ethnicity[33], identified associations with methylation at over 6,000 CpG loci[24]. We found 1341 of 4404 that passed QC in our own study (30.4%) were nominally associated with ethnicity (p < .05), which represented a highly significant (p < 2×10^-16^) enrichment. Using a Bonferroni correction for the 4404 loci tested, 126 maternal-smoking related loci were associated with ethnicity (p < 1.1×10^-5^), and 27 loci were among the 916 CpG’s reported above as associated with ethnicity [Table S7]. We also examined methylation loci from an earlier study of maternal smoking in Norwegian newborns[23] as well as studies of diesel exhaust particles[25] and exposure to violence[27]. These results are supportive of our hypothesis that environmental exposures may be responsible for the observed differences in methylation between ethnic groups and are presented in Table S8.

In an earlier study of maternal smoking in Norwegian newborns[23] that identified 26 loci associated with maternal smoking during pregnancy, 19 passed quality control (QC) in our own analysis, and the association between methylation and ethnicity was found to be nominally significant (p < 0.05)at 6 (31.6%) CpG loci. Adjusting for 19 tests (p < .0026), cg23067299 in the aryl hydrocarbon receptor repressor (*AHRR*) gene on chromosome 5 remained statistically significant [Table S8]. These results suggest that ethnic differences in methylation at loci known to be responsive to tobacco smoke exposure *in utero* may be explained in part by ethnic-specific differences in the prevalence of maternal smoking during pregnancy.

We also found that CpG loci previously reported to be associated with diesel-exhaust particle (DEP) exposure[25] were significantly enriched among the set of loci whose methylation levels varied between ethnic groups. Specifically, of the 101 CpG sites that were significantly associated with exposure to DEP and passed QC in our dataset, 31 were nominally associated with ethnicity (p < 0.05), and 5 were associated with ethnicity after adjusting for 101 comparisons (p < 0.005). Finally, we found that methylation levels at cg11218385 in the pituitary adenylate cyclase-activating polypeptide type I receptor gene (*ADCYAP1R1*), which had been associated with exposure to violence in Puerto Ricans[27] and with heavy trauma exposure in adults[28], was significantly associated with ethnicity (p = 0.02).

We also found 194 loci with a significant association between global genetic ancestry and methylation levels (after adjusting for ethnicity) at a Bonferroni corrected association p-value of less than 1.6×10^-7^ [Figure S5 and Table S9], including 48 that were associated with ethnicity in our earlier analysis. Of these significant associations, 55 were driven primarily by differences in African ancestry, 94 by differences in Native American ancestry, and 45 by differences in European ancestry. The most significant association between methylation and ancestry occurred at cg04922029 in the Duffy antigen receptor chemokine gene (DARC) on chromosome 1 (ANOVA p-value 3.1×10^-24^) [Figure S5B]. This finding was driven by a strong association between methylation level and global African ancestry; each 25 percentage point increase in African ancestry was associated with an increase in M-value of 0.98, which corresponds to an almost doubling in the ratio of methylated to unmethylated DNA at the site (95% CI 0.72 to 1.06 per 25% increase in African ancestry, p = 1.1×10^-21^). There was no significant heterogeneity in the association between genetic ancestry and methylation between Puerto Ricans and Mexicans (p-het = 0.5). Mexicans have a mean unadjusted methylation M-value 0.48 units lower than Puerto Ricans (95% CI 0.35 to 0.62 units, p = 1.1×10^-11^). However, adjusting for African ancestry accounts for the differences in methylation level between the two sub-groups (p-adjusted = 0.4), demonstrating that ethnic differences in methylation at this site are due to differences in African ancestry.

The distribution of methylation M-values at cg04922029 is tri-modal, raising the possibility that a SNP whose allele frequency differs between African and non-African populations may be driving the association. We therefore looked at the association between methylation at cg0422029 and ancestry at that locus. We found almost almost perfect correlation between methylation and African ancestry at the locus (p = 6×10^-162^) [Figure 2A]. Each African haplotype at the CpG site was associated with an increase in methylation M-value of 2.7, corresponding to a 6.5-fold increase in the ratio of methylated to unmethylated DNA per African haplotype at that locus. We then looked for SNPs within 10,000 base pairs of the CpG site that explained the admixture mapping association. We found that methylation at cg04922029 was significantly correlated with SNP rs2814778 [Figure 2B], the Duffy null mutation, 212 base pairs away; each copy of the C allele was associated with an increase in M-value of 1.5, or a 2.9-fold increase in the ratio of methylated to unmethylated DNA (p = 3.8×10^-90^) [Figure 2C].

**Figure 2:**
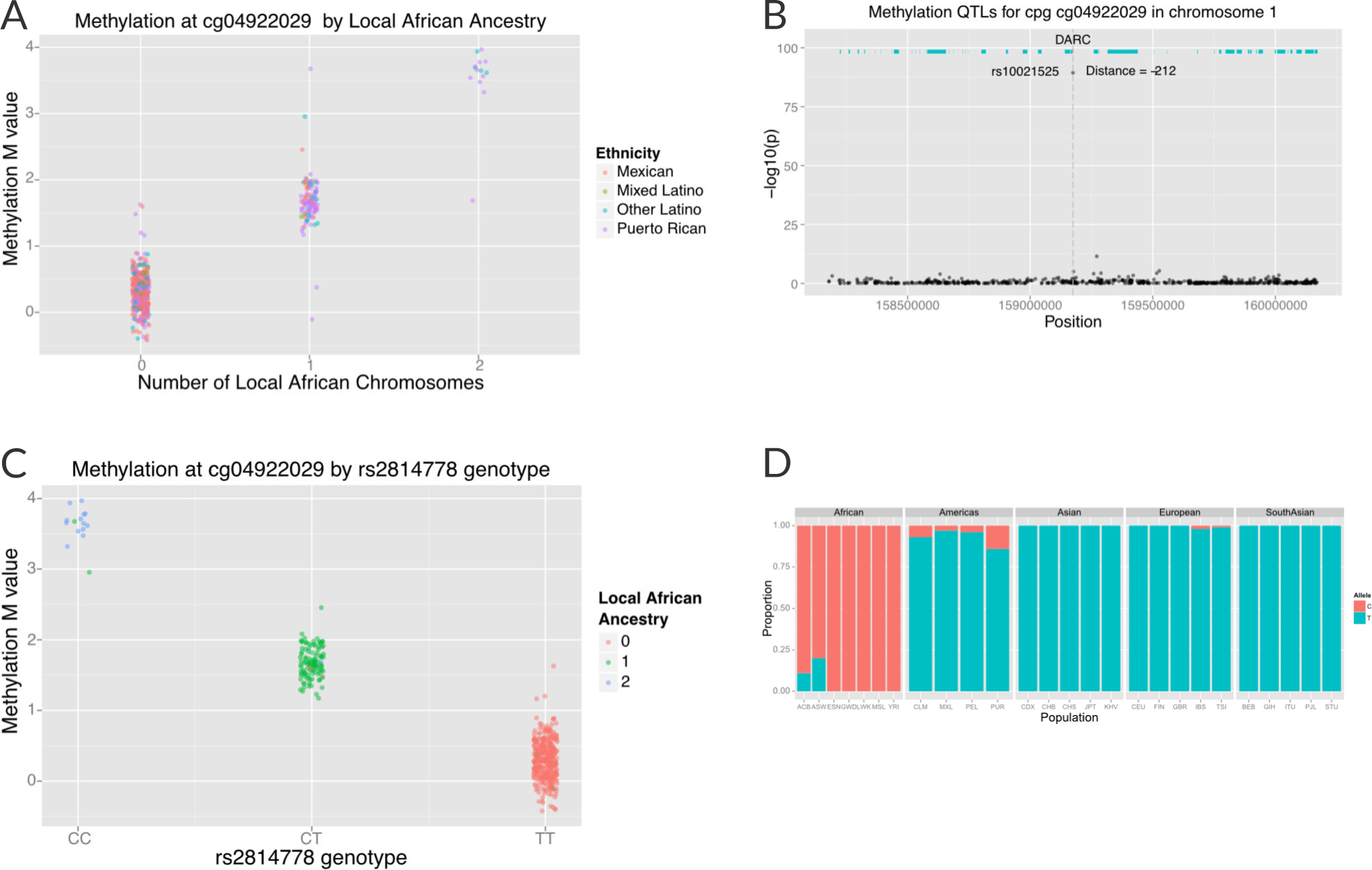
Association between local ancestry and methylation. Indicate figure parts with bold capital letters (**A**) Association between cg04922029 on the *DARC* locus and African ancestry, color coded by ethnic group. There is near perfect correlation between the two. (**B**) Association between SNPs located within 1Mb of cg04922029 and methylation levels at that CpG. (**C**) Association between rs2814778 (Duffy null) genotype and methylation at cg04922029, color coded by the number of African alleles present. There is near perfect correlation between genotype, ancestry and methylation at the locus. (**D**) Allele frequency of rs2814778 by 1000 Genomes population. The C allele is nearly ubiquitous in African populations and nearly absent outside of African populations and their descendants.

When we examined the effect of local ancestry at the other 194 CpG’s we find that a substantial proportion of the effect of global ancestry on local methylation levels is due to local ancestry acting in -cis. Among the 194 CpG sites associated with global ancestry, local ancestry at the CpG site explained a median of 10.4% (IQR 3.0% to 19.4%) of the variance in methylation, accounting for a median of 52.8% (IQR 20.3% to 84.9%) of the total variance explained jointly by local and global ancestry [Figure S6].

## Discussion

In a diverse population of Latinos, we have shown that a substantial number of loci are differentially methylated between ethnic sub-groups. While genomic ancestry can explain a portion of the association between ethnicity and methylation, factors other than shared ancestry that correlate with ethnicity, such as social, economic, cultural and environmental exposures account for the association between ethnicity and methylation at 34% (314/916) of loci.

We conclude that systematic environmental differences between ethnic subgroups likely play an important role in shaping the methylome for both individuals and populations. Loci previously associated with diverse environmental exposures such as *in utero* exposure to tobacco smoke[23,24], as well as diesel exhaust particles[25] and psychosocial stress[27] were enriched in our set of loci where methylation was associated with ethnicity. Twenty-seven of the loci associated with maternal smoking during pregnancy in a large consortium meta-analysis[24] were differentially methylated between Latino sub-groups at a genome-wide significance threshold of 1.6×10^-7^. Thus, inclusion of relevant social and environmental exposures in studies of methylation may help elucidate racial/ethnic disparities in disease prevalence, health outcomes and therapeutic response. However, in many cases, a detailed environmental exposure history is unknown, unmeasurable or poorly quantifiable, and race/ethnicity may be a useful, albeit imperfect proxy.

Our comprehensive analysis of high-density methyl- and genotyping from genomic DNA allowed us to investigate the genetic control of methylation in great detail and without the potential destabilizing effects of EBV transformation and culture in cell lines[41]. The strongest patterns of methylation are associated with cell composition in whole blood[30]. However, the specific type of Latino ethnic-subgroups (Puerto Rican, Mexican, other, or mixed) is also associated with principal coordinates of genome-wide methylation.

Our approach has some potential limitations. It is possible that fine-scale population structure (sub-continental ancestry) within European, African, and Native American populations may contribute to ethnic differences in methylation, as we had previously reported in the case of lung function[39]. However, despite the presence of additional substructure among the GALA II participants, PC’s 3-10 explained the association between ethnicity and ancestry at only 51 loci. PCs from chip-based genotypes will not capture all forms of genetic variation. Clusters of ethnicity specific rare variants of large effect or strong ethnicity-specific selective sweeps in the last 8-12 generations[37] could also give rise to methylation differences, but these are inconsistent with existing rare variant and selection analyses[42,43]. Our models of genetic ancestry assumed a linear effect of ancestry on methylation, whereas a nonlinear association or other model misspecification could have led to incomplete adjustment for genetic ancestry, and thus, led to a residual association between ethnicity and methylation. However, when we added second and third order polynomials or cubic splines to our models, we found evidence for a nonlinear association between ancestry and methylation at only 25 and 26 loci, respectively, and it did not affect the association between ethnicity and methylation. Although it is impossible to account for all types of non-linearity and non-additivity (such as gene by gene or gene by environment interaction), our analysis suggests that non-linear effects are unlikely to be significant. Since our study was geographically diverse, recruiting participants at five recruitment sites in the United States and Puerto Rico, it is possible that systematic differences associated with site of recruitment might have influenced observed methylation differences between ethnic groups. However, when we included recruitment site as a covariate, we found no significant effect on methylation independent of ethnicity.

The presence of a strong association between genetic ancestry and methylation raises the possibility that epigenetic studies can be confounded by population stratification, similar to genetic association studies, and that adjustment for either genetic ancestry or selected principal components is warranted. This possibility was first demonstrated in a previous analysis of the association between self-described race and methylation[22]. However, the study only evaluated two distinct racial groups (African Americans and Whites), while the present study demonstrates the possibility of population stratification in an admixed and heterogeneous population with participants from diverse Latino national origins. The tendency to consider Latinos as a homogenous or monolithic ethnic group makes any analysis of this population particularly challenging. Our finding of loci whose methylation patterns differed between Latino ethnic subgroups, even after adjusting for genetic ancestry, suggests that any analysis of these populations in disease-association studies without adjusting for ethnic heterogeneity is likely to result in spurious associations even after controlling for genomic ancestry.

In summary, this study provides a framework for understanding how genetic, social and environmental factors can contribute to systematic differences in methylation patterns between ethnic subgroups, even between presumably closely related populations such as Puerto Ricans and Mexicans. Methylation QTL’s whose allele frequency varies by ancestry lead to an association between local ancestry and methylation level. This, in turn, leads to systematic variation in methylation patterns by ancestry, which then contributes to ethnic differences in genome-wide patterns of methylation. However, although genetic ancestry has been used to adjust for confounding in genetic studies, and can account for much of the ethnic differences in methylation in this study, ethnic identity is associated with methylation beyond the effects of shared genetic ancestry. This is likely due to social and environmental effects captured by ethnicity. Indeed, we find that CpG sites known to be influenced by social and environmental exposures are also differentially methylated between ethnic subgroups. These findings called attention to a more complete understanding of the effect of social and environmental variables on methylation in the context of race and ethnicity to fully understanding this complex process.

Our findings have important implications for the independent and joint effects of race, ethnicity, and genetic ancestry in biomedical research and clinical practice, especially in studies conducted in diverse or admixed populations. Our conclusions may be generalizable to any population that is racially mixed such as those from South Africa, India, and Brazil, though we would encourage further study in diverse populations, and likely has implications for all studies of diverse populations. As the National Institutes of Health (NIH) embarks on a precision medicine initiative, this research underscores the importance of including diverse populations and studying factors capturing the influence of social, cultural, and environmental factors, in addition to genetic ones, upon disparities in disease and drug response.

## Materials and Methods

### Participant Recruitment

All research on human subjects was approved by the Institutional Review Board at the University of California and each of the recruitment sites (Kaiser Permanente Northern California, Children’s Hospital Oakland, Northwestern University, Children’s Memorial Hospital Chicago, Baylor College of Medicine on behalf of the Texas Children’s Hospital, VA Medical Center in Puerto Rico, the Albert Einstein College of Medicine on behalf of the Jacobi Medical Center in New York and the Western Review Board on behalf of the Centro de Neumologia Pediatrica), and all participants/parents provided age-appropriate written assent/consent. Latino children were enrolled as a part of the ongoing GALA II case-control study[33].

A total of 4,702 children (2,374 participants with asthma and 2,328 healthy controls) were recruited from five centers (Chicago, Bronx, Houston, San Francisco Bay Area, and Puerto Rico) using a combination of community- and clinic-based recruitment. Participants were eligible if they were 8-21 years of age and self-identified as a specific Latino ethnicity and had four Latino grandparents. Asthma cases were defined as participants with a history of physician diagnosed asthma and the presence of two or more symptoms of coughing, wheezing, or shortness of breath in the 2 years preceding enrollment. Participants were excluded if they reported any of the following: (1) 10 or more pack-years of smoking; (2) any smoking within 1 year of recruitment date; (3) history of lung diseases other than asthma (cases) or chronic illness (cases and controls); or (4) pregnancy in the third trimester. Further details of recruitment are described elsewhere[33]. Latino sub-ethnicity was determined by self-identification and the ethnicity of the their four grandparents. Due to small numbers, ethnicities other than Puerto Rican and Mexican were collapsed into a single category, “other Latino”. Participants whose four grandparents were of discordant ethnicity were considered to be of “mixed Latino” ethnicity.

Trained interviewers, proficient in both English and Spanish, administered questionnaires to gather baseline demographic data, as well as information on general health, asthma status, acculturation, social, and environmental exposures.

### Methylation

Genomic DNA (gDNA) was extracted from whole blood using Wizard^®^ Genomic DNA Purification Kits (Promega, Fitchburg, WI). A subset of 573 participants (311 cases with asthma and 262 healthy controls) was selected for methylation. Methylation was measured using the Infinium HumanMethylation450 BeadChip (Illumina, Inc., San Diego, CA) following the manufacturer’s instructions.

1 µg of gDNA was bisulfite-converted using the Zymo EZ DNA Methylation Kit™ (Zymo research, Irvine, CA) according to the manufacturer’s instructions. Bisulfite converted DNA was isothermally amplified overnight, enzymatically fragmented, precipitated, and re-suspended in hybridization buffer. The fragmented, re-suspended DNA samples were dispensed onto Infinitum HumanMethylation450 BeadChips and incubated overnight in an Illumina hybridization oven. Following hybridization, free DNA was washed away, and the BeadChips were extended through single nucleotide extensions with fluorescent labels. The BeadChips were imaged using an Illumina iScan system, and processed using the Illumina GenomeStudio Software.

Failed probes were identified using detection p-values using Illumina’s recommendations. Probes on sex chromosomes and those known to contain genetic polymorphisms in the probe sequence were also excluded, leaving 321,503 probes for analysis. Raw data were normalized using Illumina’s control probe scaling procedure. Beta values of methylation (ranging from 0 to 1) were converted to M-values via a logit transformation[44].

### Genotyping

Details of genotyping and quality control procedures for single nucleotide polymorphisms (SNPs) and individuals have been described elsewhere[45]. Briefly, participants were genotyped at 818,154 SNPs on the Axiom^®^ Genome-Wide LAT 1, World Array 4 (Affymetrix, Santa Clara, CA)[46]. We removed SNPs with >5% missing data and failing platform-specific SNP quality criteria (n=63,328), along with those out of Hardy-Weinberg equilibrium (n=1845; p<10-6) within their respective populations (Puerto Rican, Mexican, and other Latino), as well as non-autosomal SNPs. Subjects were filtered based on 95% call rates and sex discrepancies, identity by descent and standard Affymetrix Axiom metrics. The total number of participants passing QC was 3,804 (1,902 asthmatic cases, 1,902 healthy controls), and the total number of SNPs passing QC was 747,129. The number of participants with both methylation and genotyping data was 524.

### Ancestry and PCA analysis

GALA II participants were combined with ancestral data from 1000 Genomes European (CEU) and African (YRI) populations and 71 Native American (NAM) samples genotyped on the Axiom^®^ Genome-Wide LAT 1 array. A final sample of 568,037 autosomal SNPs with relevant ancestral data was used to estimate local and global ancestry. Global ancestry was estimated using the program ADMIXTURE[47], with a three population model. Local ancestry at all positions across the genome was estimated using the program LAMP-LD[48], assuming three ancestral populations.

Principal components for the genetic data were determined using the program EIGENSTRAT[40].

### Statistical analysis

Unless otherwise noted, all regression models were adjusted for case status, age, sex, estimated cell counts, and plate and position. To account for possible heterogeneity in the cell type makeup of whole blood we inferred white cell counts using the method by Houseman et al[35]. Indicator variables were used to code categorical variables with more than two categories, such as ethnicity. In these cases, a nested analysis of variance (ANOVA) was used to compare models with and without the variables to obtain an omnibus p-value for the association between the categorical variable and the outcome. For analyses of dependent beta-distributed variables (such as African, European, and Native American ancestries), or cell proportion, k-1 variables were included in the analysis, and a nested analysis of variance (ANOVA) was used to compare models with and without the variables to obtain an k-1 degree of freedom omnibus p-value for the association between predictor (such as ancestry) and the outcome variable.

The Bonferroni method was used to adjust for multiple comparisons. For methylome-wide associations, the significance threshold was adjusted for 321,503 probes, resulting in a Bonferroni threshold of 1.6×10^-7^. Analyses were performed using R version 3.2.1 (The R Foundation for Statistical Computing)[49] and the Bioconductor package version 2.13.

Multidimensional scaling of the logit transformed methylation data (M-values) was performed by first calculating the Euclidian distance matrix between each pair of individuals and then calculating the first 10 principal coordinates of the data [Figure S2A]. We performed both a simple correlation analysis of these principal coordinates to demographic factors (age, sex, ethnicity), estimated cell counts and technical factors (batch, plate, and position) to identify factors that correlated with global methylation patterns [see Figure S2B]. In addition, we performed a multiple regression analysis of methylation principal coordinates by ethnicity and ancestry, adjusting for case status, age, sex, estimated cell counts, and plate and position [Table S1].

We also sought to establish the extent to which global differences in methylation between Puerto Ricans and Mexicans could be explained by differences in ancestry between the two groups. We estimated the proportion of the ethnicity association that was mediated by genomic ancestry using the R package “mediation”[36] for methylation principal coordinates, which demonstrated a significant association with ethnicity.

We also sought to correlate ethnicity and methylation at a locus-specific level. We thus performed a linear regression between methylation at each CpG site and self-reported ethnicity (Mexican, Puerto Rican, Mixed Latino, and Other Latino), followed by a three degree of freedom analysis of variance to determine the overall effect of ethnicity on methylation We repeated the analysis excluding the 16 participants that were self-described as “Mixed Latino”, and tested for non-linearity in two ways: by adding second and third order polynomials to the model, and by adding a 3-degree of freedom cubic spline and comparing models with the non-linear terms to those without using a nested ANOVA. At loci where there was evidence for non-linearity, we tested whether ethnicity remained associated with methylation after adjusting for ancestry as well as the deviations from linearity. Finally, we tested for the presence of population sub-structure beyond that conveyed through ancestry by adding the genetic principal components 3-10 (PCs 1 and 2 were co-linear with ancestry with a correlation coefficient R2 > 0.998) and comparing models with those PCs to those without. At loci where there was evidence for association between PC’s 3-10 and methylation, we tested whether ethnicity remained associated with methylation after adjusting for ancestry as well as the PC’s 3-10.

We calculated the proportion of variance in methylation explained by ethnicity and genomic ancestry at each site where ethnicity was significantly associated with methylation. To do this, we fit a model that included both ethnicity and global ancestry as well as the confounders described above and calculated the proportion of variance explained by multiplying the ratio of the variance between predictors (ethnicity and genomic ancestry) and outcome (methylation) by the square of the effect magnitude (ß).

We also examined whether differences in methylation patterns by ethnicity could be associated with known loci that had previously been reported to vary based on common environmental exposures, including maternal smoking during pregnancy[23], diesel exhaust particles (DEP)[25], and exposure to violence[27]. We have previously shown that exposure to these common environmental exposures or similar exposures varied by ethnicity within our own GALA II study populations[33,50,51].

In addition, we examined the association between global ancestry and methylation across all CpG loci using a two-degree of freedom likelihood ratio test as well as by examining the association between individual ancestral components (African, European, and Native American) and methylation at each CpG site. At each site where methylation was significantly associated with genomic ancestry proportions, we determined the relative effect of global ancestry (θ, theta) and local ancestry (γ, gamma) in a joint model by calculating the proportion of variance explained as above.

To determine whether ancestry associations with methylation were due to variation in local ancestry, we correlated local ancestry at each CpG site with methylation at the site. Because ancestry LD is much stronger than genotypic LD, it is possible to accurately interpolate ancestry at each CpG site based on the ancestry estimated at the nearest SNPs[45,52]. Measures of locus-specific ancestry were correlated with local methylation using linear regression. We performed a two-degree of freedom analysis of variance test evaluating the overall effect of all three ancestries as well as single-ancestry associations comparing methylation at a given locus with the number of African, European and Native American chromosomes at that CpG site.

## Acknowledgements

The authors acknowledge the families and patients for their participation and thank the numerous health care providers and community clinics for their support and participation in GALA II. In particular, the authors thank study coordinator Sandra Salazar; the recruiters who obtained the data: Duanny Alva, MD, Gaby Ayala-Rodriguez, Lisa Caine, Elizabeth Castellanos, Jaime Colon, Denise DeJesus, Blanca Lopez, Brenda Lopez, MD, Louis Martos, Vivian Medina, Juana Olivo, Mario Peralta, Esther Pomares, MD, Jihan Quraishi, Johanna Rodriguez, Shahdad Saeedi, Dean Soto, Ana Taveras. We also thank Sasha Gusev for helpful discussion.

Computations in this manuscript were performed using the UCSF Biostatistics High Performance Computing System.

## References

1. Risch N, Burchard E, Ziv E, Tang H. Categorization of humans in biomedical research: genes, race and disease. Genome Biology. BioMed Central; 2002;3: comment2007.

2. Cooper RS, Kaufman JS, Ward R. Race and genomics. N Engl J Med. 2003;348: 1166–1170. doi:10.1056/NEJMsb022863

3. Yudell M, Roberts D, Desalle R, Tishkoff S. SCIENCE and SOCIETY. Taking race out of human genetics. Science. American Association for the Advancement of Science; 2016;351: 564–565. doi:10.1126/science.aac4951

4. Hankinson JL, Odencrantz JR, Fedan KB. Spirometric reference values from a sample of the general U.S. population. Am J Respir Crit Care Med. American Thoracic Society New York, NY; 1999;159: 179–187. doi:10.1164/ajrccm.159.1.9712108

5. Quanjer PH, Stanojevic S, Cole TJ, Baur X, Hall GL, Culver BH, et al. Multi-ethnic reference values for spirometry for the 3–95-yr age range: the global lung function 2012 equations. European Respiratory Journal. European Respiratory Society; 2012;40: 1324–1343. doi:10.1183/09031936.00080312

6. Borrell LN. Racial identity among Hispanics: implications for health and well-being. Am J Public Health. 2005;95: 379–381. doi:10.2105/AJPH.2004.058172

7. Kumar R, Seibold MA, Aldrich MC, Williams LK, Reiner AP, Colangelo L, et al. Genetic Ancestry in Lung-Function Predictions. N Engl J Med. 2010;363: 321–330. doi:10.1056/NEJMoa0907897

8. Udler MS, Nadkarni GN, Belbin G, Lotay V, Wyatt C, Gottesman O, et al. Effect of Genetic African Ancestry on eGFR and Kidney Disease. J Am Soc Nephrol. American Society of Nephrology; 2015;26: 1682–1692. doi:10.1681/ASN.2014050474

9. Nalls MA, Wilson JG, Patterson NJ, Tandon A, Zmuda JM, Huntsman S, et al. Admixture Mapping of White Cell Count: Genetic Locus Responsible for Lower White Blood Cell Count in the Health ABC and Jackson Heart Studies. The American Journal of Human Genetics. 2008;82: 81–87. doi:10.1016/j.ajhg.2007.09.003

10. Nguyen AB, Moser R, Chou WY. Race and health profiles in the United States: an examination of the social gradient through the 2009 CHIS adult survey. Public Health. 2014;128: 1076–1086. doi:10.1016/j.puhe.2014.10.003

11. Evans GW, Kantrowitz E. Socioeconomic Status and Health: The Potential Role of Environmental Risk Exposure. Annual Review of Public Health. Annual Reviews 4139 El Camino Way, P.O. Box 10139, Palo Alto, CA 94303–0139, USA; 2002;23: 303–331. doi:10.1146/annurev.publhealth.23.112001.112349

12. Smith ZD, Meissner A. DNA methylation: roles in mammalian development. Nat Rev Genet. 2013;14: 204–220. doi:10.1038/nrg3354

13. Kulis M, Esteller M. DNA methylation and cancer. Adv Genet. Elsevier; 2010;70: 27–56. doi:10.1016/B978-0-12-380866-0.60002-2

14. Udali S, Guarini P, Moruzzi S, Choi S-W, Friso S. Cardiovascular epigenetics: from DNA methylation to microRNAs. Mol Aspects Med. 2013;34: 883–901. doi:10.1016/j.mam.2012.08.001

15. Kato N, Loh M, Takeuchi F, Verweij N, Wang X, Zhang W, et al. Trans-ancestry genome-wide association study identifies 12 genetic loci influencing blood pressure and implicates a role for DNA methylation. Nat Genet. Nature Publishing Group; 2015. doi:10.1038/ng.3405

16. Bell CG, Finer S, Lindgren CM, Wilson GA, Rakyan VK, Teschendorff AE, et al. Integrated genetic and epigenetic analysis identifies haplotype-specific methylation in the FTO type 2 diabetes and obesity susceptibility locus. Sorensen TIA, editor. PLoS ONE. Public Library of Science; 2010;5: e14040. doi:10.1371/journal.pone.0014040

17. Chambers JC, Loh M, Lehne B, Drong A, Kriebel J, Motta V, et al. Epigenome-wide association of DNA methylation markers in peripheral blood from Indian Asians and Europeans with incident type 2 diabetes: a nested case-control study. Lancet Diabetes Endocrinol. 2015;3: 526–534. doi:10.1016/S2213-8587(15)00127-8

18. Liu Y, Aryee MJ, Padyukov L, Fallin MD, Hesselberg E, Runarsson A, et al. Epigenome-wide association data implicate DNA methylation as an intermediary of genetic risk in rheumatoid arthritis. Nat Biotechnol. Nature Publishing Group; 2013;31: 142–147. doi:10.1038/nbt.2487

19. Lardenoije R, Iatrou A, Kenis G, Kompotis K, Steinbusch HWM, Mastroeni D, et al. The epigenetics of aging and neurodegeneration. Prog Neurobiol. 2015;131: 21–64. doi:10.1016/j.pneurobio.2015.05.002

20. Bell JT, Pai AA, Pickrell JK, Gaffney DJ, Pique-Regi R, Degner JF, et al. DNA methylation patterns associate with genetic and gene expression variation in HapMap cell lines. Genome Biology. BioMed Central Ltd; 2011;12: R10. doi:10.1186/gb-2011-12-1-r10

21. Feil R, Fraga MF. Epigenetics and the environment: emerging patterns and implications. Nat Rev Genet. Nature Publishing Group; 2011;13: 97–109. doi:10.1038/nrg3142

22. Barfield RT, Almli LM, Kilaru V, Smith AK, Mercer KB, Duncan R, et al. Accounting for population stratification in DNA methylation studies. Genet Epidemiol. 2014;38: 231–241. doi:10.1002/gepi.21789

23. Joubert BR, Håberg SE, Nilsen RM, Wang X, Vollset SE, Murphy SK, et al. 450K epigenome-wide scan identifies differential DNA methylation in newborns related to maternal smoking during pregnancy. Environ Health Perspect. 2012;120: 1425–1431. doi:10.1289/ehp.1205412

24. Joubert BR, Felix JF, Yousefi P, Bakulski KM, Just AC, Breton C, et al. DNA Methylation in Newborns and Maternal Smoking in Pregnancy: Genome-wide Consortium Meta-analysis. Am J Hum Genet. 2016;98: 680–696. doi:10.1016/j.ajhg.2016.02.019

25. Jiang R, Jones MJ, Sava F, Kobor MS, Carlsten C. Short-term diesel exhaust inhalation in a controlled human crossover study is associated with changes in DNA methylation of circulating mononuclear cells in asthmatics. Part Fibre Toxicol. 2014;11: 71. doi:10.1186/s12989-014-0071-3

26. Ho S-M, Johnson A, Tarapore P, Janakiram V, Zhang X, Leung Y-K. Environmental epigenetics and its implication on disease risk and health outcomes. ILAR J. Oxford University Press; 2012;53: 289–305. doi:10.1093/ilar.53.3-4.289

27. Chen W, Boutaoui N, Brehm JM, Han Y-Y, Schmitz C, Cressley A, et al. ADCYAP1R1 and asthma in Puerto Rican children. Am J Respir Crit Care Med. 2013;187: 584–588. doi:10.1164/rccm.201210-1789OC

28. Ressler KJ, Mercer KB, Bradley B, Jovanovic T, Mahan A, Kerley K, et al. Post-traumatic stress disorder is associated with PACAP and the PAC1 receptor. Nature. 2011;470: 492–497. doi:10.1038/nature09856

29. van der Knaap LJ, Riese H, Hudziak JJ, Verbiest MMPJ, Verhulst FC, Oldehinkel AJ, et al. Glucocorticoid receptor gene (NR3C1) methylation following stressful events between birth and adolescence. The TRAILS study. Transl Psychiatry. Nature Publishing Group; 2014;4: e381. doi:10.1038/tp.2014.22

30. Lam LL, Emberly E, Fraser HB, Neumann SM, Chen E, Miller GE, et al. Factors underlying variable DNA methylation in a human community cohort. Proceedings of the National Academy of Sciences. National Acad Sciences; 2012;109 Suppl 2: 17253–17260. doi:10.1073/pnas.1121249109

31. Borghol N, Suderman M, McArdle W, Racine A, Hallett M, Pembrey M, et al. Associations with early-life socio-economic position in adult DNA methylation. International Journal of Epidemiology. Oxford University Press; 2012;41: 62–74. doi:10.1093/ije/dyr147

32. Vidal AC, Benjamin Neelon SE, Liu Y, Tuli AM, Fuemmeler BF, Hoyo C, et al. Maternal stress, preterm birth, and DNA methylation at imprint regulatory sequences in humans. Genet Epigenet. Libertas Academica; 2014;6: 37–44. doi:10.4137/GEG.S18067

33. Oh SS, Tcheurekdjian H, Roth LA, Nguyen EA, Sen S, Galanter JM, et al. Effect of secondhand smoke on asthma control among black and Latino children. J Allergy Clin Immunol. 2012;129: 1478–1483.e7. doi:10.1016/j.jaci.2012.03.017

34. Michels KB, Binder AM, Dedeurwaerder S, Epstein CB, Greally JM, Gut I, et al. Recommendations for the design and analysis of epigenome-wide association studies. Nat Methods. 2013;10: 949–955. doi:10.1038/nmeth.2632

35. Houseman EA, Accomando WP, Koestler DC, Christensen BC, Marsit CJ, Nelson HH, et al. DNA methylation arrays as surrogate measures of cell mixture distribution. BMC Bioinformatics. BioMed Central Ltd; 2012;13: 86. doi:10.1186/1471-2105-13-86

36. Tingley D, Yamamoto T, Hirose K, Keele L, Imai K. mediation: R package for causal mediation analysis. UCLA Statistics/American Statistical Association. UCLA Statistics/American Statistical Association; 2014.

37. Galanter JM, Fernandez-Lopez JC, Gignoux CR, Barnholtz-Sloan J, Fernandez-Rozadilla C, Via M, et al. Development of a panel of genome-wide ancestry informative markers to study admixture throughout the Americas. Gibson G, editor. PLoS Genet. Public Library of Science; 2012;8: e1002554. doi:10.1371/journal.pgen.1002554

38. Bryc K, Velez C, Karafet T, Moreno-Estrada A, Reynolds A, Auton A, et al. Colloquium paper: genome-wide patterns of population structure and admixture among Hispanic/Latino populations. Proceedings of the National Academy of Sciences. 2010;107 Suppl 2: 8954–8961. doi:10.1073/pnas.0914618107

39. Moreno-Estrada A, Gignoux CR, Fernandez-Lopez JC, Zakharia F, Sikora M, Contreras AV, et al. Human genetics. The genetics of Mexico recapitulates Native American substructure and affects biomedical traits. Science. American Association for the Advancement of Science; 2014;344: 1280–1285. doi:10.1126/science.1251688

40. Patterson N, Price AL, Reich D. Population Structure and Eigenanalysis. PLoS Genet. Public Library of Science; 2006;2: e190. doi:10.1371/journal.pgen.0020190

41. Grafodatskaya D, Choufani S, Ferreira JC, Butcher DT, Lou Y, Zhao C, et al. EBV transformation and cell culturing destabilizes DNA methylation in human lymphoblastoid cell lines. Genomics. 2010;95: 73–83. doi:10.1016/j.ygeno.2009.12.001

42. Hernandez RD, Kelley JL, Elyashiv E, Melton SC, Auton A, McVean G, et al. Classic selective sweeps were rare in recent human evolution. Science. American Association for the Advancement of Science; 2011;331: 920–924. doi:10.1126/science.1198878

43. Tang H, Choudhry S, Mei R, Morgan M, Rodríguez-Cintrón W, Burchard EG, et al. Recent Genetic Selection in the Ancestral Admixture of Puerto Ricans. The American Journal of Human Genetics. 2007;81: 626–633. doi:10.1086/520769

44. Du P, Zhang X, Huang C-C, Jafari N, Kibbe WA, Hou L, et al. Comparison of Beta-value and M-value methods for quantifying methylation levels by microarray analysis. BMC Bioinformatics. BioMed Central Ltd; 2010;11: 587. doi:10.1186/1471-2105-11-587

45. Galanter JM, Gignoux CR, Torgerson DG, Roth LA, Eng C, Oh SS, et al. Genome-wide association study and admixture mapping identify different asthma-associated loci in Latinos: the Genes-environments & Admixture in Latino Americans study. J Allergy Clin Immunol. 2014;134: 295–305. doi:10.1016/j.jaci.2013.08.055

46. Hoffmann TJ, Zhan Y, Kvale MN, Hesselson SE, Gollub J, Iribarren C, et al. Design and coverage of high throughput genotyping arrays optimized for individuals of East Asian, African American, and Latino race/ethnicity using imputation and a novel hybrid SNP selection algorithm. Genomics. Elsevier Inc; 2011;: 1–9. doi:10.1016/j.ygeno.2011.08.007

47. Alexander DH, Novembre J, Lange K. Fast model-based estimation of ancestry in unrelated individuals. Genome Research. Cold Spring Harbor Lab; 2009;19: 1655–1664. doi:10.1101/gr.094052.109

48. Baran Y, Pasaniuc B, Sankararaman S, Torgerson DG, Gignoux C, Eng C, et al. Fast and accurate inference of local ancestry in Latino populations. Bioinformatics. 2012;28: 1359–1367. doi:10.1093/bioinformatics/bts144

49. Team RC. R: A language and environment for statistical computing [Internet]. 3rd ed. Vienna. Available: http://www.R-project.org/

50. Nishimura KK, Galanter JM, Roth LA, Oh SS, Thakur N, Nguyen EA, et al. Early-life air pollution and asthma risk in minority children. The GALA II and SAGE II studies. Am J Respir Crit Care Med. 2013;188: 309–318. doi:10.1164/rccm.201302-0264OC

51. Thakur N, Oh SS, Nguyen EA, Martin M, Roth LA, Galanter J, et al. Socioeconomic status and childhood asthma in urban minority youths. The GALA II and SAGE II studies. Am J Respir Crit Care Med. 2013;188: 1202–1209. doi:10.1164/rccm.201306-1016OC

52. Rosenberg NA, Huang L, Jewett EM, Szpiech ZA, Jankovic I, Boehnke M. Genome-wide association studies in diverse populations. Nat Rev Genet. Nature Publishing Group; 2010;11: 356–366. doi:10.1038/nrg

